# Modulation of tactile detection threshold with rhythmic somatosensory entrainment

**DOI:** 10.1101/695692

**Authors:** Michel J. Wälti, Marc Bächinger, Nicole Wenderoth

**Affiliations:** Neural Control of Movement Lab, Department of Health Sciences and Technology, ETH Zurich, Switzerland; Neuroscience Center Zurich (ZNZ), University and ETH Zurich, Zurich, Switzerland

**Author notes:** Correspondence: Michel J. Wälti. **Author contributions** MJW and MB designed research; MJW performed research and analyzed data; MJW, MB, and NW wrote the paper. **Funding** This research was funded by the Swiss National Science Foundation (No. 320030_175616).

**Keywords:** steady-state evoked potentials, entrainment, somatosensory system, neural oscillations, vibrotactile stimulation, tactile detection

## Abstract

Ongoing neural activity in human somatosensory cortex has a strong impact on the detectability of weak tactile stimuli. Recent studies suggest that brain oscillations, which determine the state of excitability of a cortical area, play a crucial role in this process. Mainly two frequency bands have been reported to be involved in conscious sensory perception: alpha (8 – 12 Hz) and beta (15 – 30 Hz). In addition to correlative findings, more recent studies investigated causality by measuring the extent to which directly modulating brain oscillations affects sensory perception. While most of these studies use transcranial alternating current stimulation (tACS), rhythmic sensory stimulation has been suggested as a simple and safe alternative to entrain ongoing neural activity. However, convincing findings demonstrating the modulation of neural signals and related behavioral function are scarce.

Here, we investigated whether rhythmically induced brain oscillations by means of vibrotactile stimulation (i.e. sensory entrainment) modulate tactile detection. In line with previous findings, we show in trials without sensory entrainment that preceding alpha power and phase-angles in beta oscillations predict the detection rate of a weak tactile stimulus. Further, we reveal a masking effect induced by sensory entrainment stimulation resulting in higher perception thresholds. Intriguingly, we find that the masking effect is modulated by the strength of neural entrainment resulting from 20 Hz stimulation. Our data provide evidence for the possibility to modulate sensory processing with rhythmic sensory stimulation. However, in light of the induced masking effects, the feasibility of this entrainment method to modulate human behavior remains questionable.

## Introduction

Whether a weak stimulus is consciously perceived or not depends not only on the characteristics of the stimulus, but also on ongoing neuronal processes. A variety of recent studies suggest that part of the variability in perception can be explained by the state of excitability of a cortical area as reflected by the specific signature of the underlying brain oscillations (Hanslmayr, Gross, Klimesch, & Shapiro, 2011; Iemi, Chaumon, Crouzet, & Busch, 2017; Jensen & Mazaheri, 2010; Klimesch, Sauseng, & Hanslmayr, 2007; Lange, Oostenveld, & Fries, 2013; Limbach & Corballis, 2016).

Among studies investigating oscillatory activity prior to weak stimuli in perceptual detection tasks, mainly two frequency bands have been reported to be involved in conscious sensory perception: alpha (8 – 12 Hz) and beta (15 – 30 Hz). Alpha band power in occipital areas has been shown in many studies to be correlated with visual perception (Hanslmayr et al., 2007; Iemi et al., 2017; Mathewson, Gratton, Fabiani, Beck, & Ro, 2009; Rajagovindan & Ding, 2011; Romei, Gross, & Thut, 2010; Thut, Nietzel, Brandt, & Pascual-Leone, 2006; van Dijk, Schoffelen, Oostenveld, & Jensen, 2008). In the somatosensory system, not only alpha, but also beta band power and phase characteristics have been reported to be related to tactile processing of near-threshold stimuli (Ai & Ro, 2014; Baumgarten, Schnitzler, & Lange, 2015; Linkenkaer-Hansen, Nikulin, Palva, Ilmoniemi, & Palva, 2004; Palva, Linkenkaer-Hansen, Naatanen, & Palva, 2005; Schubert, Haufe, Blankenburg, Villringer, & Curio, 2009; Zhang & Ding, 2010). Linkenkaer-Hansen et al. (2004) found participants’ detection rate to a near-threshold stimulus to be improved with intermediate levels of preceding oscillatory power in the somatosensory system for both, alpha and beta. More recent studies found similar effects of alpha and beta, however, most report a more linear relationship between oscillatory power and sensory detection. For example, Frey et al. (2016), which had participants perform a near-threshold detection task, found that low alpha and beta power in the contralateral somatosensory region predicted whether a weak stimulus was detected (Frey et al., 2016). While most of these findings are based on electroencephalography (EEG) or magnetoencephalography (MEG) recordings and are correlative in nature, more recent studies investigated the causality of this effect by directly modulating brain oscillations to reveal their effect on sensory perception.

Transcranial alternating current stimulation (tACS) is a common method to alter neuronal activity and has been reported to affect power, phase and frequency of ongoing brain oscillations (e.g. Bächinger et al., 2017; Cecere, Rees, & Romei, 2015; Zaehle, Rach, & Herrmann, 2010). In the visual domain, 10 Hz tACS over occipital areas has been shown to modulate visual detection performance in a phase-dependent manner (Helfrich et al., 2014). In the somatosensory domain, tACS at participants’ individual alpha frequency (IAF) has been shown to modulate tactile detection: Gundlach et al. (2016) stimulated participants’ bilateral primary somatosensory cortices with tACS while a tactile detection task was performed. Although no overall effect of tACS on detection threshold was reported, variations in performance were found regarding the phase of ongoing stimulation in relation to the onset of the tactile stimulus (Gundlach et al., 2016). In addition, it has been shown that alpha power in the somatosensory system decreased after tACS (Gundlach, Muller, Nierhaus, Villringer, & Sehm, 2017) and that this decrease would result in an increase of reporting rates in a tactile detection task (increase in hit rates but also in false alarm rates) (Craddock, Poliakoff, El-Deredy, Klepousniotou, & Lloyd, 2017). Although tACS represents a widely used method for modulating brain oscillations, online interactions between the externally applied electric fields and the ongoing endogenous brain rhythms are still poorly understood. This is at least partly because electrical artifacts induced by tACS cause a rhythmic contamination that is several magnitudes larger than the recorded EEG signals (Thut, Schyns, & Gross, 2011).

An alternative method to entrain brain oscillations is rhythmic sensory stimulation, which induces steady-state evoked potentials (SSEPs) in corresponding cortical areas following the temporal frequency of the driving stimulus (Regan, 1977). SSEPs have the advantage that they can be measured simultaneously with EEG and have been documented in the visual (steady-state visually evoked potentials, SSVEPs), the auditory (auditory steady-state responses, ASSRs) and the somatosensory (steady-state somatosensory evoked potentials, SSSEPs) systems (for a review, see Vialatte, Maurice, Dauwels, & Cichocki, 2010). The underlying mechanism of SSEPs, however, is still debated (see Zoefel, Ten Oever, & Sack, 2018). Although some studies found evidence for entrained brain oscillations by revealing an interaction between stimulation and ongoing neuronal activity (Notbohm, Kurths, & Herrmann, 2016; Schwab et al., 2006; Wälti, Bächinger, Ruddy, & Wenderoth, 2019), findings that show an established behavioral modulation as a consequence of rhythmically entrained brain oscillations are scarce (see Haegens & Zion Golumbic, 2018).

The present study aims to investigate whether rhythmically induced brain oscillations via vibrotactile stimulation reveal a behavioral effect on tactile detection thresholds and/or detection rates, thus demonstrating behavioral evidence for neural entrainment as a result of rhythmic sensory stimulation. In line with previous findings, we focus on alpha (10 Hz) and beta (20 Hz) stimulation frequencies, applied to the thumb (dominant hand). To uncover entrainment effects on a neurophysiological level, we measure EEG throughout the experiment and apply jittered stimulation signals with an average frequency of 20 Hz as an additional control condition.

## Materials and methods

### Participants

40 healthy right-handed participants were recruited and tested in this study. 5 participants were excluded from the analysis due to technical problems during the EEG recording or because they did not follow the instructions, resulting in a final sample of 35 participants (female: 22; age: M ± SD = 24.0 ± 4.1). The study protocol was approved by the local ethics committee and was conducted in accordance with the Declaration of Helsinki. Participants signed an informed consent and were informed about the procedure of the experiment, but were kept naïve with regard to the hypothesis of the study.

### Design and procedure

Participants were comfortably seated in a dark sound-attenuated room, approximately 80 – 120 cm away from a computer screen (27-inch). EEG data were recorded throughout the whole experiment (see below). Participants were allowed to comfortably rest their right arm on a pillow placed on a desk next to them to ensure a relaxed position of the stimulated fingers for the duration of the experiment.

Participants were instructed to indicate with a button press whether a weak tactile stimulus (tactile burst; sine wave at 205 Hz for 250 ms) administered to the tip of the index finger of their right hand was perceived or not. In some trials, they received a vibrotactile stimulation (sensory entrainment) of 2 s duration to their thumb of the right hand prior to performing the tactile detection task for stimuli applied to the index finger (Figure 1). In total 6 entrainment conditions were used covering a range of different vibrotactile frequencies and intensities. The entrainment conditions consisted of 10 Hz, 20 Hz and jittered vibration (with an average of 20 on-off cycles per second), each at a low and high stimulation intensity. Vibrotactile stimulation signals were generated as previously reported (Wälti et al., 2019). The behavioral effect of vibrotactile entrainment was compared to a control condition (no entraining stimulation) by determining individual detection thresholds using a staircase procedure (Quest algorithm from PsychToolbox; Watson & Pelli, 1983). The detection threshold was determined over 25 trials converging on 82 % correct answers (as predicted by the Weibull psychometric function; see Watson & Pelli, 1983) and revealed the minimal intensity of the tactile stimulation to be perceived by the participant. Immediately after determination of the threshold estimation, participants were asked to perform 10 additional trials of the perceptual detection task, each trial using tactile stimulation at the previously estimated threshold. For each of the 7 conditions (6 entrainment conditions and 1 control condition), this procedure of threshold estimation followed by trials at estimated threshold was performed once per block in randomized order. In total, the experiment consisted of 3 blocks, resulting in 3 estimated thresholds and 30 (3 × 10) trials at individual perception threshold for each condition.

**Figure 1.**
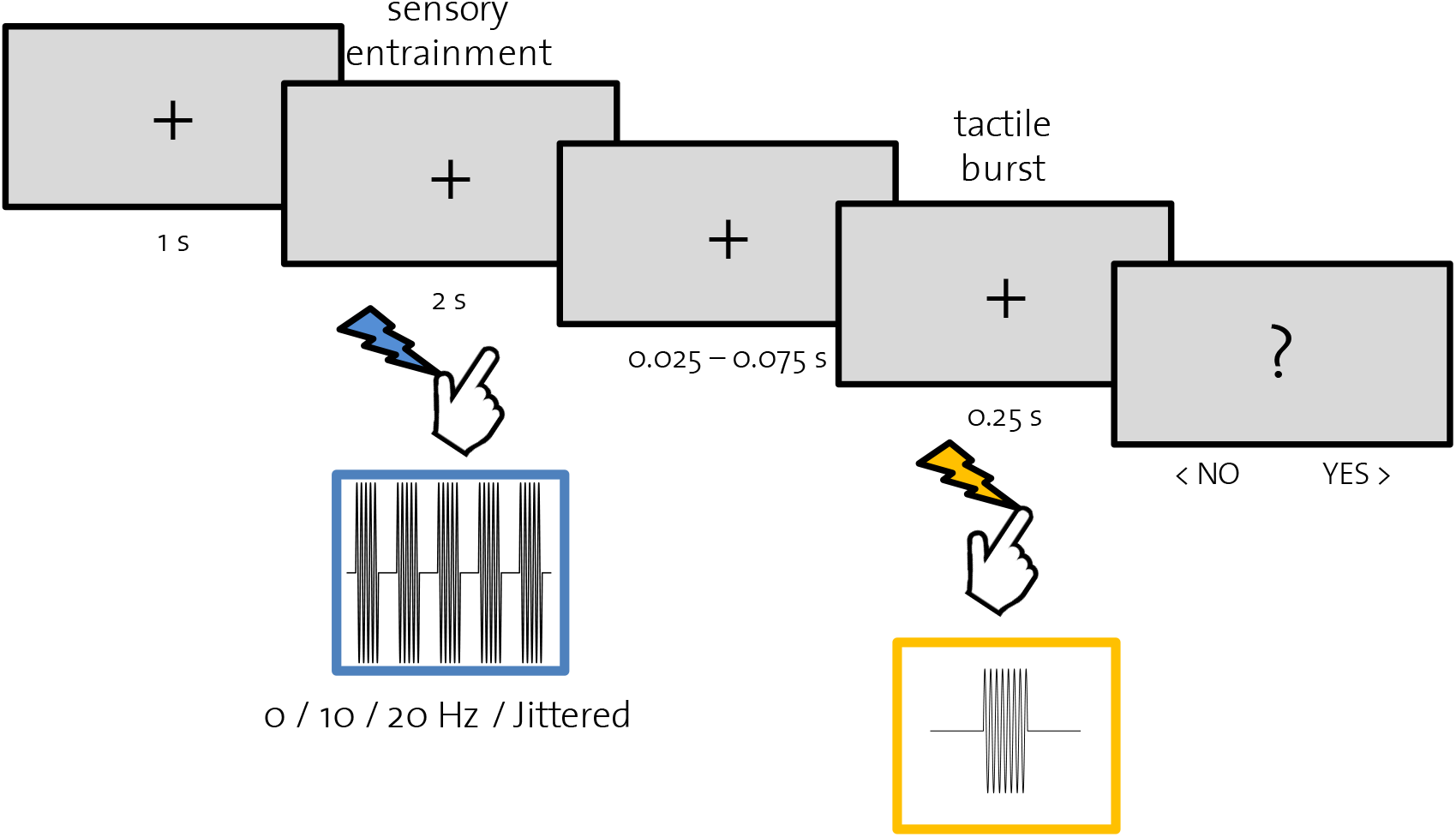
Experimental procedure. Perceptual detection task at index finger preceded by vibrotactile stimulation at thumb. Stimulation was administered to the right hand and responses were given using the left hand. The procedure was the same for threshold estimation and subsequent trials at individual perception threshold.

### Behavioral data analysis

In order to compare individual estimated thresholds across conditions, for each block we subtracted the estimated threshold in the control condition from values of the 6 entrainment conditions. Then, z-scores were calculated from log-transformed threshold differences (∆ estimated threshold). To determine statistical differences between stimulation conditions, we used R (R Core Team, 2018) and the lme4 package (Bates, Mächler, Bolker, & Walker, 2015) to perform a linear mixed effects analysis. The linear mixed effects model was used to account for variance arising from individual differences and systematic differences across trials (Baayen, Davidson, & Bates, 2008). To this end, subjects and trial number were included in the model as random factors. Fixed effects represent the influence of the stimulation conditions on the dependent variable (here: ∆ estimated threshold).

### EEG data acquisition and preprocessing

EEG data were acquired at 1000 Hz using a 64-channel Hydrocel Geodesic EEG System (Electrical Geodesic Inc., USA), referenced to Cz (vertex), with an online Notch filter (50 Hz) and high-pass filter at 0.3 Hz. Impedances were kept below 50 kΩ.

EEG data were preprocessed and analyzed offline. Data were cleaned (detection and interpolation of bad electrodes), band-pass filtered (0.5 – 30 Hz) and further processed using independent component analysis (ICA). Artifact components (ICs) were automatically detected and removed from the data with a custom built toolbox (see Liu, Ganzetti, Wenderoth, & Mantini, 2018). Finally, EEG data were average re-referenced.

### EEG data analysis

To determine electrophysiological correlates of sensory entrainment and perceptual performance, we determined for each participant the electrodes over the contralateral somatosensory region showing the strongest responses to alpha (10 Hz) and beta (20 Hz) stimulation, both delivered at high intensity. For this, data from each channel from high alpha and high beta stimulation conditions were epoched separately (−0.5 s to +2.0 s regarding stimulation onset).

To estimate power across time and frequencies, data from each channel and trial were convolved with a family of complex Morlet wavelets spanning 6 – 24 Hz in 20 steps with wavelet cycles increasing logarithmically between 4 and 10 cycles as a function of frequency. Alpha (9.5 – 10.5 Hz) and beta (19.5 – 20.5 Hz) power were obtained by squaring the absolute value averaged over time points and trials. Further, power values were normalized to baseline (−0.5 s to −0.25 s) by converting to the decibel scale (Cohen, 2014). For subsequent analyses involving 10 Hz entrainment conditions, EEG data at each subject’s strongest alpha electrode were used. Likewise, analyses involving 20 Hz and jittered conditions were based on EEG data at the individual strongest beta electrode.

To test the hypotheses regarding prestimulus alpha and beta power, and phase angle determining subsequent sensory detection performance, data from the control condition were analyzed. For this, we epoched data around the onset of tactile bursts (−0.5 s to +1.0 s) and derived power and phase values from a time-frequency analysis as described above. Pre-stimulus alpha and beta power was calculated as averaged power across 500 ms preceding stimulus onset for each trial separately. Further, for each trial, we derived phase angles at stimulation onset at 10 and 20 Hz. Pre-stimulus power values were further divided in three bins with an equal number of trials. For each power bin, detection rates were assessed. To determine phase angle effects on detection performance across levels of low and high frequency-specific power, pre-stimulus power values were divided in two bins (low and high power) and each bin was further divided in peaks and troughs, with regard to the phase angle value of the signal at stimulation onset. If phase angle values exceeded π/2, they were identified as peak trials, else, as trough trials. Again, detection rates were calculated and compared across the power and phase angle bins.

To show general EEG effects of sensory entrainment, alpha and beta power (baseline corrected, see before) were calculated and averaged across trials for each stimulation condition separately.

In order to further test effects of sensory entrainment on perceptual performance, we analyzed 10 and 20 Hz stimulation conditions and compared low and high stimulation intensities. Trials from each condition were divided into trough and peak trials using the same procedure as described above.

To determine the level of entrainment, we measured changes in intersite phase-clustering (ISPC), which is a measure of phase-based connectivity between two time-series signals (Cohen, 2014). ISPC was calculated between the averaged time-series data of each stimulation condition and the corresponding stimulation signal (see Wälti et al., 2019).

A repeated-measures ANOVA with Bonferroni-corrected post-hoc pairwise comparisons and paired t-tests in SPSS Version 25 (IBM, USA) were used for further statistical analyses.

## Results

The current study was designed to measure the effect of sensory entrainment through vibrotactile stimulation of the thumb on subsequent tactile detection performance of the index finger. Our goal was to investigate the importance of alpha and beta power and phase angles on sensory perception and to further evaluate the entrainment of ongoing brain oscillations via rhythmic sensory stimulation as a possible mechanism for modulating behavior.

### Effects of preceding alpha power and beta phase angles on detection rates

First, we analyzed control condition data (no sensory entrainment) in order to replicate previous work that demonstrated that preceding alpha and beta power, and phase angles are associated with sensory detection performance. Comparison of detection rates across three levels of preceding alpha power revealed an optimum at an intermediate level as reported in previous studies (e.g. Linkenkaer-Hansen et al., 2004; Figure 2 A). A repeated-measures ANOVA revealed a significant effect of power level (F(2,33) = 4.456, p = 0.019) and post-hoc pairwise comparison showed a significant difference between intermediate and high levels (p = 0.039). No significant difference was found between low and intermediate, and low and high levels of alpha power. No effects on detection rates were found between phase angles in low and high alpha trials (2×2 ANOVA with factors power and phase; main effects: power: F(1,34) = 0.622, p = 0.436, phase: F(1,34) = 0.166, p = 0.686; interaction: F(1,34) = 0.009, p = 0.923; Figure 2 B). Preceding beta revealed a linear decrease of detection rate with increasing power (Figure 2 C). Although not reaching significance (F(2,33) = 0.451, p = 0.641) this is in line with previous reports (e.g. Jones et al., 2010). Further, detection rates in high beta trials revealed a significant dependence on phase angle values (t(34) = 2.264, p = 0.030). As previously reported, sensory detection was enhanced during troughs compared to peaks (Figure 2 D). This confirms previous results showing that beta oscillations define perceptual cycles in the somatosensory cortex (Baumgarten et al., 2015).

**Figure 2.**
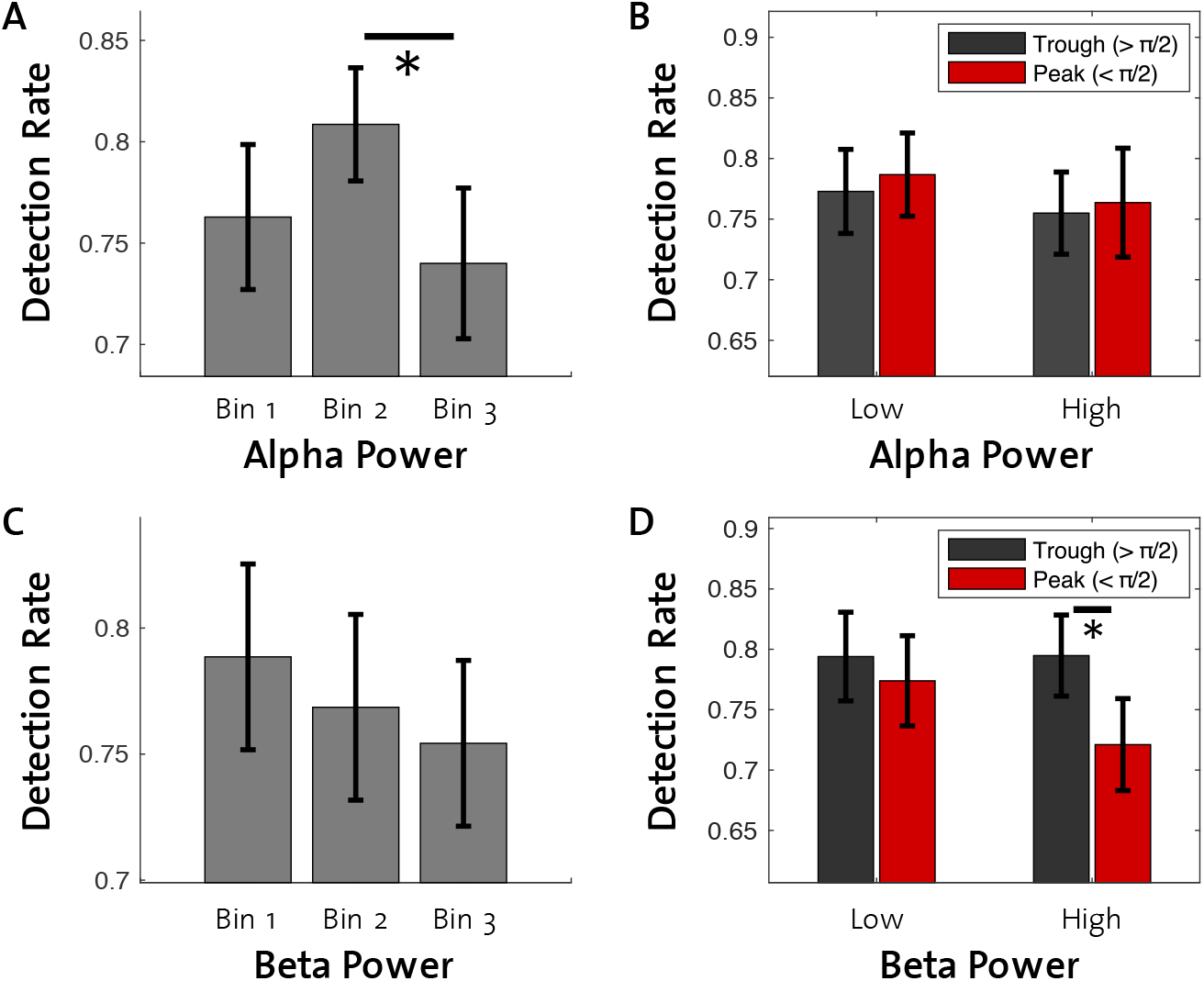
Effects of preceding alpha and beta on detection rates in the control condition. Detection rates were measured as mean number of detected stimuli at estimated thresholds in the control condition (no sensory entrainment). A: Preceding alpha power reveals an optimum at an intermediate level for tactile detection. B: No phase-dependent effect on detection was found in low and high alpha power trials. C: On average, detection rates decrease with increasing preceding beta power (not significant). D: High beta power trials reveal a phase-dependent effect on detection. Tactile burst presentation at troughs of the beta phase lead to higher detection rates compared to presentation during peaks. (* = p < 0.05, error bars = standard error of mean, SEM)

### Sensory entrainment reveals effects on beta but not alpha power

In order to determine whether our stimulation protocol reveals typical characteristics of steady-state responses over the somatosensory area across alpha and beta frequencies, we looked at 10 and 20 Hz power during the stimulation phase of each of the 6 entrainment conditions. Figure 3 depicts topographical power during high-intensity 10 and 20 Hz stimulation (Figure 3 A, C) and the resulting power spectrum derived from the strongest electrode, marked as a red dot (Figure 3 B, D). Highest power values were found over contralateral somatosensory areas for both stimulation conditions. Note that these data were derived from EEG data averaged across all subjects and trials (‘supersubject’), hence showing stronger effects compared to individual data. Figure 4 depicts alpha and beta power relative to the control condition for all entrainment conditions. To our surprise, statistical analysis revealed no effect of stimulation condition on alpha power (F(5,30) = 0.241, p = 0.941). Beta power on the other hand, was clearly increased by high intensity 20 Hz entrainment stimulation (F(5,30) = 10.647, p < 0.001), revealing highly significant differences compared to the other conditions in pairwise post-hoc tests (p < 0.001 for all comparisons).

**Figure 3.**
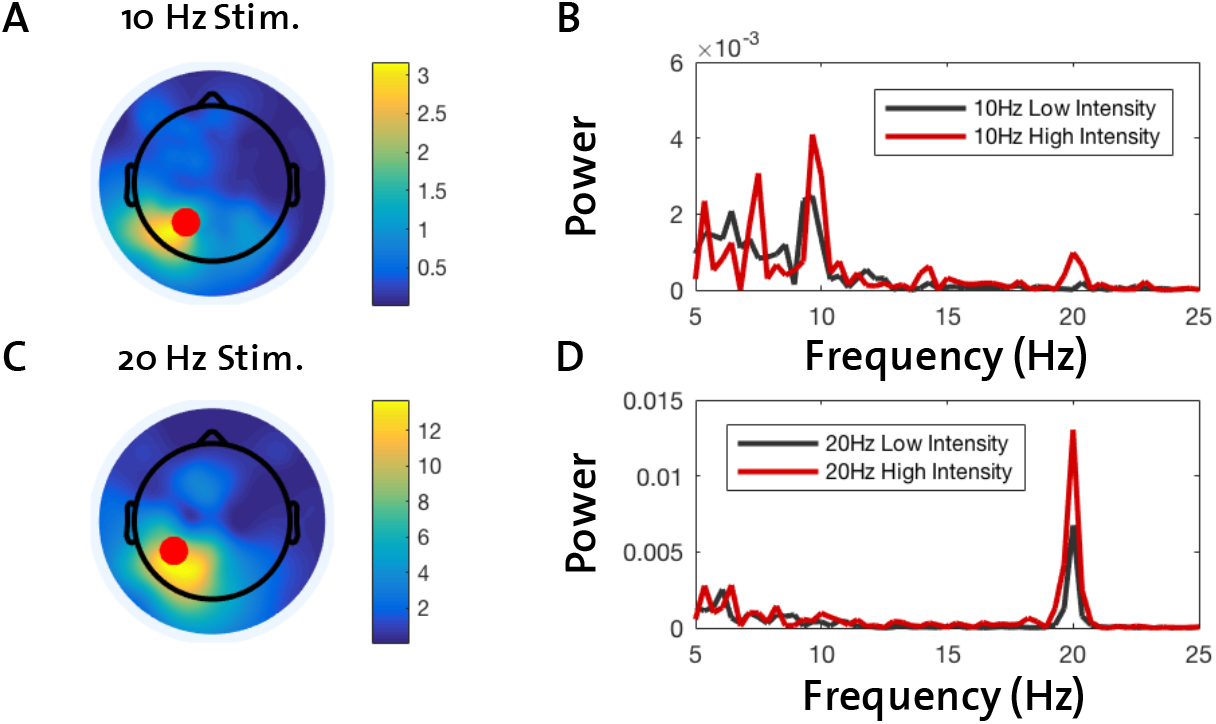
Averaged effects of high-intensity 10 and 20 Hz stimulation. A: Topographical depiction of electrode power at 10 Hz reveals strongest response to high 10 Hz stimulation over left somatosensory regions (contralateral to vibrotactile stimulation). Red dot indicates electrode with strongest 10 Hz response. B: Power spectrum derived from the strongest electrode from A in response to high 10 Hz stimulation. C: Electrode power at 20 Hz in response to high 20 Hz stimulation. Red dot indicates electrode with strongest 20 Hz response. C: Equivalent to B for high 20 Hz stimulation.

**Figure 4.**
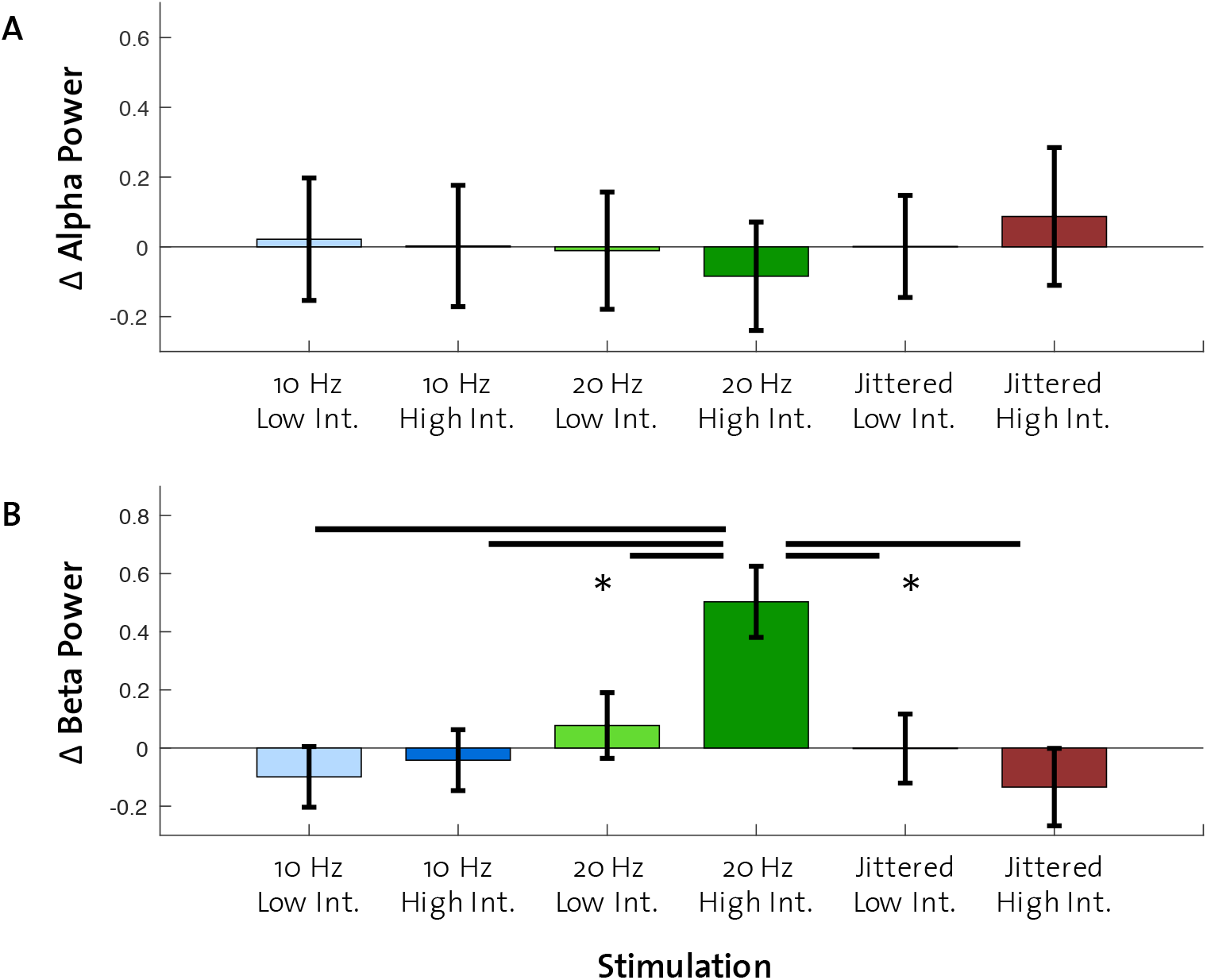
Effects of all stimulation conditions on alpha and beta power. A: Alpha power changes compared to control reveal no effect of stimulation. B: High intensity 20 Hz stimulation leads to strongest increase (compared to control) in beta power across stimulation conditions. (* = p < 0.001, error bars = SEM)

### No phase-specific effects on detection rate from low and high stimulation intensity

So far, our analyses of the control condition have revealed that the detection rate was influenced by preceding alpha power and the phase of high power beta oscillations. Further, we found that strongest beta power increases (compared to control) resulted from high-intensity 20 Hz stimulation. Consequently, we argue that a phase-dependent effect on detection rate should be found in high-intensity 20 Hz stimulation conditions. However, as depicted in Figure 5, none of our rhythmic stimulation conditions resulted in a difference in detection rates with regard to their phase angle at the onset of the tactile burst. We compared detection rates from trough and peak trials for each of the stimulation conditions separately using paired t-tests. Note that a direct comparison of detection rates between low and high stimulation intensities (main effect of intensity) would be problematic, as detection rates were derived for different estimated thresholds for each condition.

**Figure 5.**
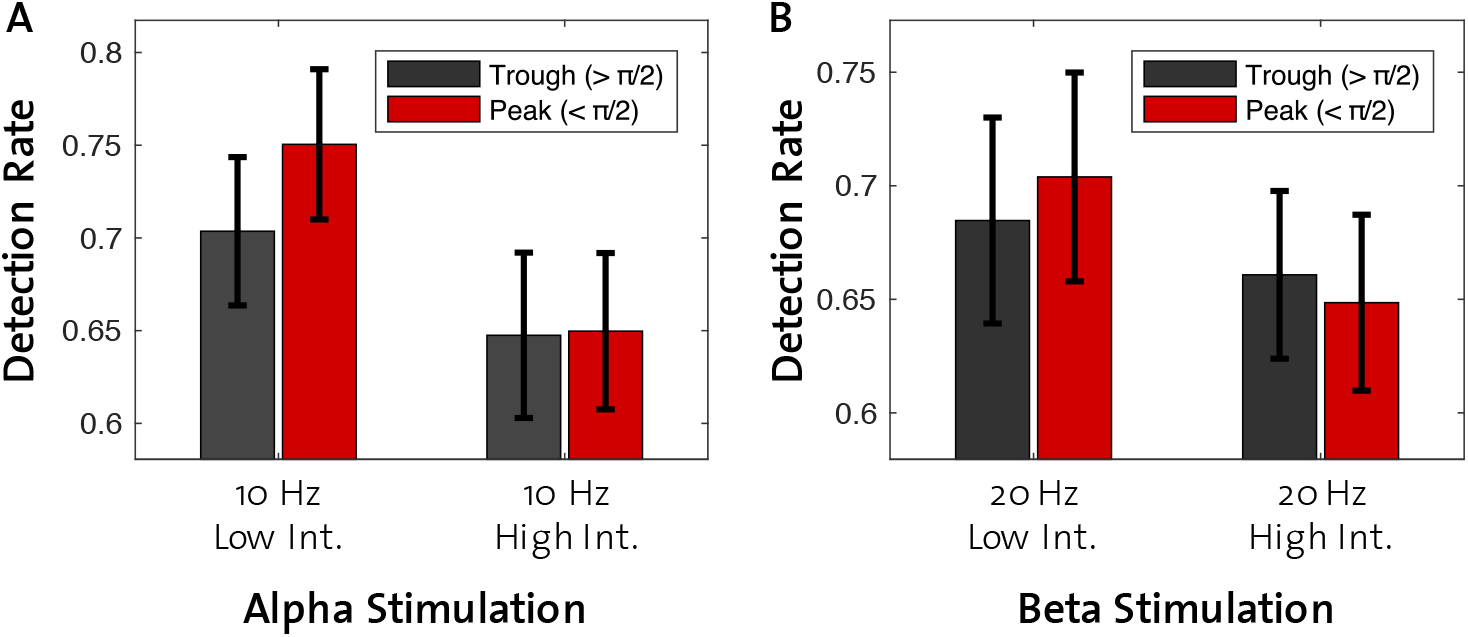
Alpha and beta stimulation conditions do not result in phase-dependent effects on detection rate. A: Trials with stimulus presentation at troughs and peaks do not differ with regard to detection rate in low- and high-intensity 10 Hz stimulation conditions. B: Equivalent of A for 20 Hz stimulation conditions. (error bars = SEM)

### Masking effect of stimulation intensity on estimated detection threshold

Sensory entrainment affected perceptual thresholds (Figure 6) as confirmed by a main effect of stimulation intensity (p < 0.001). Stimulation frequency and the interaction between frequency and intensity revealed no significant differences (10 Hz vs. 20 Hz: p = 0.294; 10 Hz vs. jittered: p = 0.532; 20 Hz vs. jittered: p = 0.937; 10 Hz low:high vs. 20 Hz low:high: p = 0.319; 10 Hz low:high vs. jittered low:high: p = 0.639; 20 Hz low:high vs. jittered low:high: p = 0.830; Figure 6). The main effect of stimulation intensity can be interpreted as a masking effect caused by the preceding vibrotactile stimulation. High intensity stimulation resulted in higher estimated detection thresholds compared to low intensity stimulation, independent of rhythmicity of the stimulation (rhythmic vs. jittered) or the stimulation frequency (10 Hz vs. 20 Hz). As depicted in Figure 6, preceding stimulation (sensory entrainment) increased perceptual thresholds in comparison to control, in each condition, with stronger masking effects for high intensity stimulation. Masking refers to the phenomenon in which one stimulus decreases the detectability of another (e.g. Gilson, 1969). Here, the effect is caused by forward masking, because the to-be-detected stimulus (tactile burst) was preceded by a masking stimulus (sensory entrainment).

**Figure 6.**
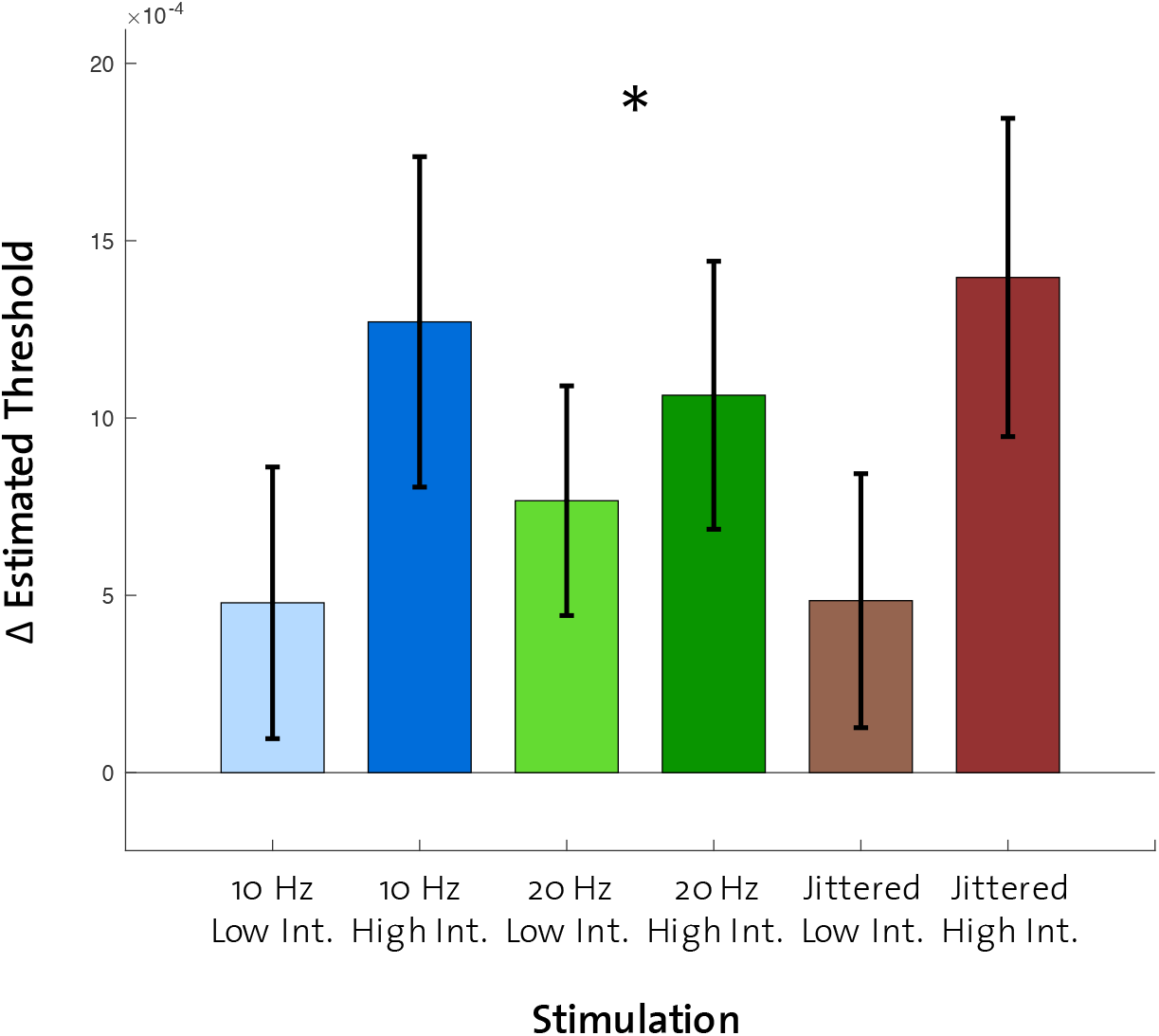
Estimated perceptual thresholds derived from all stimulation conditions in comparison to control. Increased thresholds resulting from higher stimulation intensities across all stimulation conditions can be interpreted as stronger masking effects. (* = p < 0.001, error bars = SEM)

### Beta stimulation results in highest phase-coupling (ISPC)

Next, we tested whether individual differences in the magnitude of phase-locking between stimulation signal and EEG response revealed changes in perceptual performance. As measure of phase-locking, we measured inter-site phase clustering (ISPC) between EEG signals and entrainment signals for each condition (see Wälti et al., 2019).

In line with previous findings, we found ISPC to be highest for the 20 Hz stimulation frequency and with high intensity compared to low intensity (2×2 ANOVA: main effects: frequency: F(2,33) = 42.216, p < 0.001, intensity: F(1,34) = 24.674, p < 0.001; interaction: F(2,33) = 2.550, p = 0.093; Figure 8).

**Figure 7.**
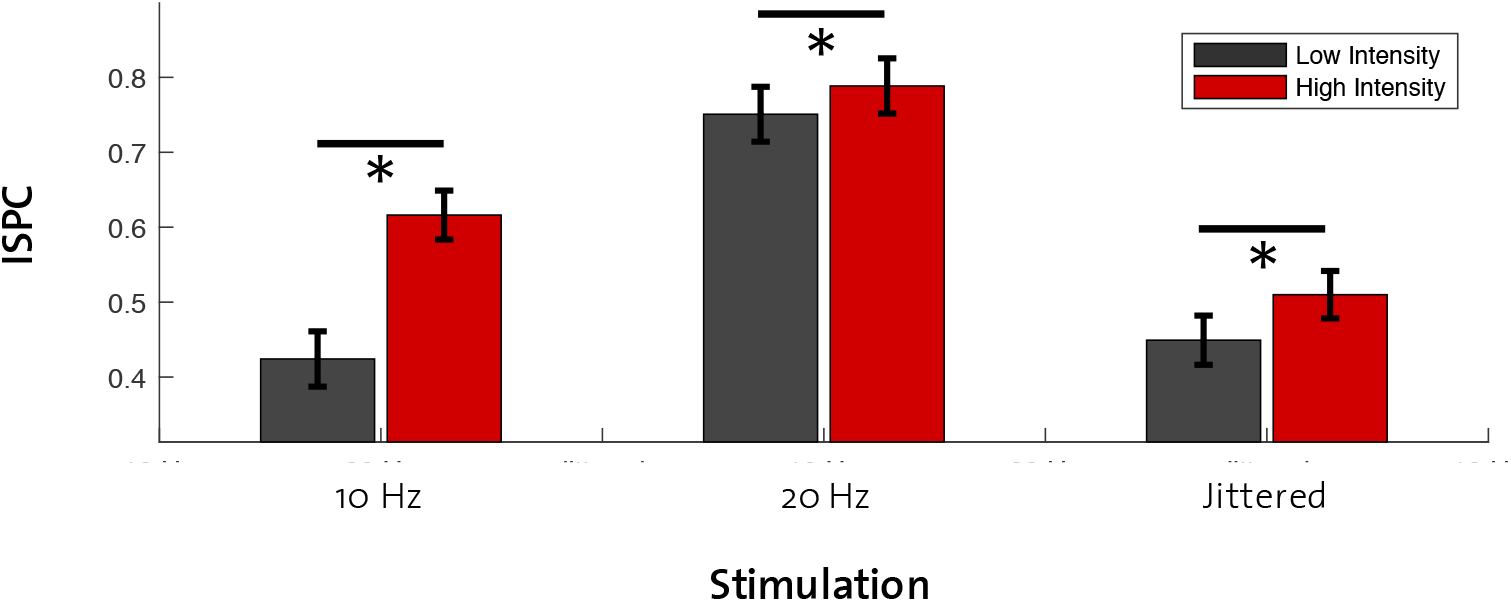
ISPC across stimulation conditions. Phase-coupling between sensory entrainment signal and resulting EEG time-series is most pronounced for 20 Hz stimulation. Further, in all conditions, high intensity resulted in higher ISPC. (* = p < 0.05, error bars = SEM)

### ISPC increase between low and high beta stimulation correlates with difference in perceptual thresholds

In a final step, we calculated ISPC differences (∆ ISPC) and tested whether a stronger gain of entrainment from low-to high-intensity stimulation also results in stronger behavioral effects. Differences in estimated perceptual thresholds between low and high stimulation intensities (high 10 Hz – low 10 Hz; high 20 Hz – low 20 Hz; high jittered – low jittered) were used as a behavioral measure (∆ TH). Correlational analyses revealed a positive relationship between ∆ ISPC and ∆ TH for the 20 Hz stimulation (r = 0.42, p = 0.016; Figure 8 B). No significant correlations were found for 10 Hz (Figure 8 A) and jittered stimulation (Figure 8 C) conditions (10 Hz: r = 0.06, p = 0.727; jittered: r = - 0.06, p = 0.731).

To further investigate the positive correlation between ∆ ISPC and ∆ TH in the 20 Hz stimulation condition, we divided the subjects into two equally sized groups using a median split based on ∆ ISPC. Figure 8 D shows that the differential behavior of the two groups was driven by the response to high-intensity 20 Hz stimulation: individuals with high ∆ ISPC had a significantly higher detection threshold for high intensity entrainment than individuals with low ∆ ISPC (paired t-tests: high intensity: p = 0.036). No such effect was found for low-intensity 20 Hz stimulation (paired t-tests: low intensity: p = 0.554).

**Figure 8.**
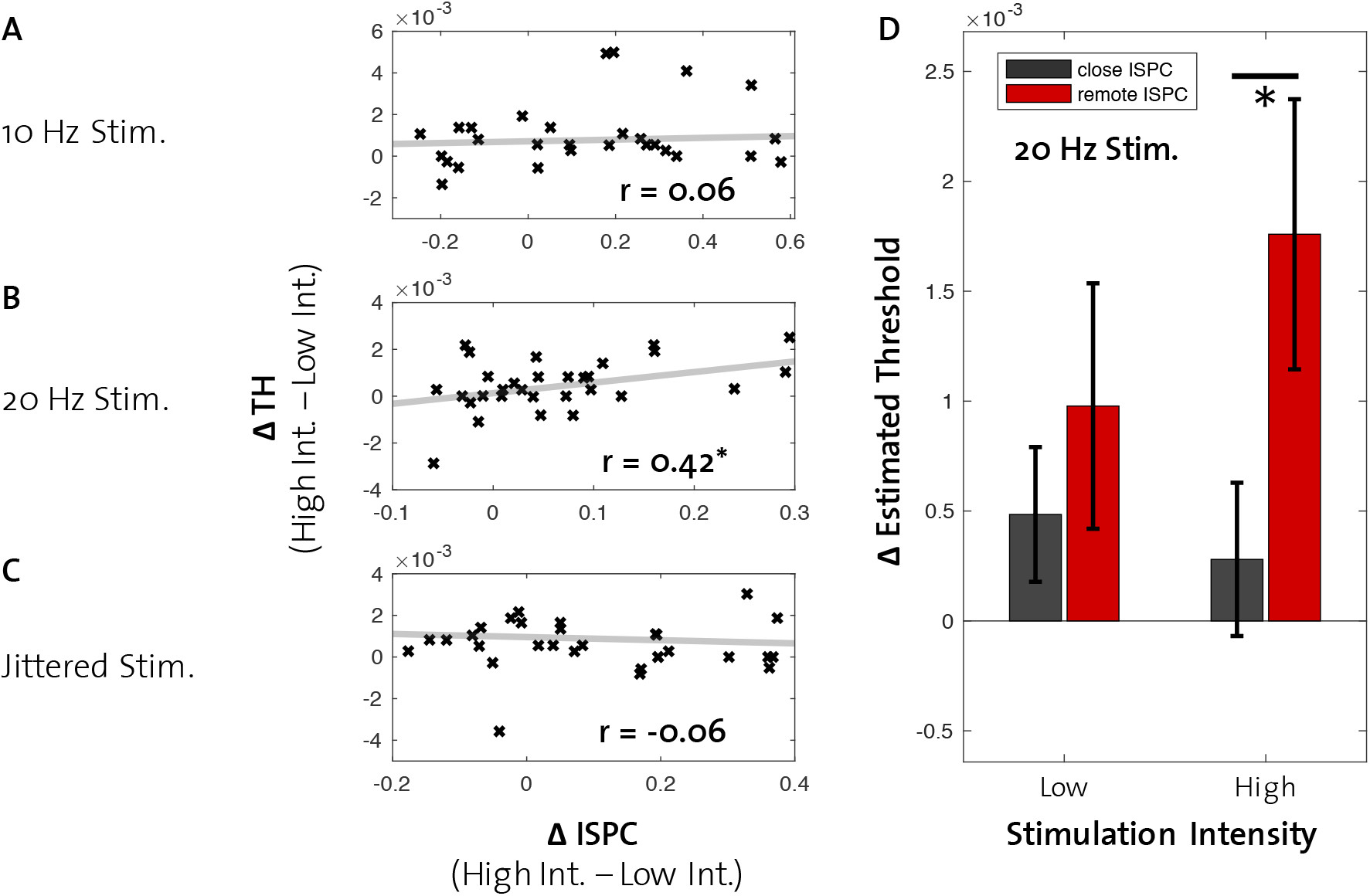
Correlations between ∆ ISPC and ∆ TH. ∆ ISPC was calculated as difference between low- and high-intensity stimulation for 10 Hz, 20 Hz and jittered stimulation separately. ∆ TH refers to estimated thresholds in low-intensity conditions subtracted from estimated thresholds in high-intensity conditions. A: No correlation was found for the 10 Hz stimulation frequency. B: A positive correlation was found for 20 Hz stimulation. C: Jittered stimulation revealed no significant correlation. D: Further investigation of the positive correlation in the 20 Hz condition revealed that participants with closer ISPC values differed from participants with more remote ISPC values in the high-intensity entrainment condition, whereas ∆ estimated threshold from low-intensity entrainment did not differ between the two groups. (* = p < 0.05, error bars = SEM)

## Discussion

In the present study, we investigated the effects of sensory entrainment on detection of a tactile stimulus. For this, we applied various rhythmic vibrotactile stimulations to the thumb, followed by a short to-be-detected stimulus to the index finger. For each condition, we estimated the detection threshold and subsequently estimated the detection rate of stimuli presented at the estimated threshold. Building on previous findings, we focused on alpha (10 Hz) and beta (20 Hz) entrainment frequencies, hypothesizing decreasing detection performance with increasing power in both frequency bands (Baumgarten et al., 2015; Ergenoglu et al., 2004; Mazaheri & Jensen, 2010; Romei et al., 2008; Schubert et al., 2009; van Dijk et al., 2008; Weisz et al., 2014). Further, phase-dependent effects in alpha and beta have been reported previously, revealing the importance of ongoing neural oscillations in somatosensory cortex (Baumgarten et al., 2015; Mathewson et al., 2011; Mazaheri & Jensen, 2010). Improved detectability of a weak stimulus can be predicted if stimulus onset is at the trough of the ongoing oscillation, in comparison to onset at the peak.

### Mixed relationships between preceding alpha and beta, and detection rate

We first tested the validity of our hypotheses by inspecting preceding alpha and beta power in the control condition (no sensory entrainment). In line with previous reports, we found an intermediate level of preceding alpha power to result in the highest detectability of a weak stimulus (administered at estimated threshold). In contrast, preceding beta revealed a more linear relationship (i.e. higher beta power results in lower detection performance). However, statistically, different levels of beta power did not result in variations in detection performance. In order to reveal phase-dependent effects of alpha and beta oscillations on detection rate, we further divided preceding neural activity in both frequency bands into low- and high-power trials. Both power levels were then further divided into two groups of trials dependent on the phase angle of the ongoing oscillation at the onset of the tactile stimulus (troughs and peaks). While no phase-dependent effect was found in both alpha power levels, high beta power trials revealed greater detectability of a stimulus onset at troughs compared to peaks.

In general, although not conclusively, our results show that preceding power in the alpha and beta band is related to the detectability of a weak sensory stimulus. In line with our results, one of the first studies investigating preceding neural activity as a predictor of detectability of a tactile stimulus revealed similar findings. Linkenkaer-Hansen et al. (2004) revealed an inverted-U relationship between alpha and beta power and subsequent detection of a weak electrical stimulus, suggesting an intermediate level of neural activity facilitating sensory processing (Linkenkaer-Hansen et al., 2004). While others have found similar effects (Ai & Ro, 2014; Zhang & Ding, 2010), some studies found a more linear relationship. Especially in the alpha band, it has been shown that correctly perceived weak tactile stimuli are preceded by low power in somatosensory cortical areas (Schubert et al., 2009; Weisz et al., 2014). The finding of a linear relationship is usually interpreted according to the notion that alpha reflects the local cortical excitability (i.e. increased alpha reflects functional inhibition) (Harvey et al., 2013; Iemi et al., 2017). On the other hand, an inverted-U relationship has been explained by either sensorimotor processing using intrinsic stochastic resonance in which the ongoing activity plays the role of an intrinsic noise source (see Linkenkaer-Hansen et al., 2004), or, a failure to reach neuronal firing thresholds in a neural system with low levels of spontaneous activity, and reduced sensory-evoked responses in response to excessive levels of spontaneous oscillations (see Zhang & Ding, 2010). In conclusion, the relationship between alpha and beta power, and sensory perception, especially in the somatosensory system, is not fully understood. Our data further suggest that such a potential brain-behavior relationship is most likely weak, resulting in borderline statistical effects and slight inconsistencies across studies.

The finding of a phase-dependent effect on detection rate in high beta power trials is in line with previous work reporting that sensory stimulation is discretely processed, and that the underlying perceptual cycles are determined by ongoing alpha and beta cycles (Baumgarten et al., 2015; Klimesch et al., 2007; Mathewson et al., 2009; Mathewson et al., 2011; Mazaheri & Jensen, 2010). These findings suggest a link between oscillatory phase angle at stimulus onset and detection performance. Alpha and beta activity is thought to produce pulses of inhibition that appear during the peaks of the oscillatory cycle. As a result, neural firing and sensory perception is reduced at these specific phases. In line with the present finding, Ai and Ro (2013) reported that preceding oscillatory power (in their study: alpha) and phase are interrelated with respect to their effect on tactile perception. They found that when alpha power was high and the stimulus was presented during a trough, detection rates were significantly higher than for those presented during a peak (Ai & Ro, 2014). However, a similar interpretation of our results is difficult, because we did not find a power-specific effect of beta on detection rate and therefore cannot argue that increases in detection rate resulting from different power levels would overwrite phase-specific effects.

With regard to the mixed alpha and beta results in the present study, it needs to be acknowledged that our experiment was not primarily designed to reveal effects of ongoing neural activity on tactile detection, but rather to compare perceptual changes resulting from sensory entrainment to a control condition. Thus, only a small fraction of trials was used to determine effects in the control condition, which was further reduced by dividing trials into different power levels.

### 20 Hz entrainment increases beta power, but does not result in phase-effects on detection rate

The purpose of the sensory entrainment used in this study was to modulate neural processing of perceptual information by changing ongoing brain oscillations in the alpha (10 Hz) and beta (20 Hz) band. In order to detect effects of our entrainment signals on EEG activity, we measured frequency-specific power changes during stimulation in comparison to control. We found that 20 Hz high-intensity stimulation resulted in a clear beta power increase compared to control and other entrainment conditions. To our surprise, no effect on alpha power was detected during 10 Hz high-intensity stimulation. One explanation for this lack of alpha response to our entrainment signal could be an already increased level of alpha power during the control condition. A relaxed state is known to be associated with increased alpha-band activity (Adrian & Yamagiwa, 1935). In addition, sensory stimulation results in a desynchronization in the alpha band (Pfurtscheller & Lopes da Silva, 1999), thus diminishing the enhancement effect of sensory entrainment on alpha power. Further, previous studies have shown that vibrotactile stimulation results in the strongest responses with frequencies in the beta range (Müller, Neuper, & Pfurtscheller, 2001; Snyder, 1992; Tobimatsu, Zhang, & Kato, 1999; Tobimatsu, Zhang, Suga, & Kato, 2000; Wälti et al., 2019). These findings have been interpreted as a resonance mechanism in the somatosensory system, which represents the frequency band in which sensory information is processed optimally (Hutcheon & Yarom, 2000). Such a resonance mechanism could be responsible for the lack of an effect of 10 Hz stimulation in our study. ISPC values derived across all stimulation conditions confirms a resonance-like mechanism. In line with previous findings, we found phase-coupling between the sensory entrainment signal and resulting EEG time-series to be pronounced for 20 Hz stimulation.

Taking together our findings of a phase-dependent effect on detection rate in trials with high beta power and the strong beta power differences resulting from low- and high-intensity 20 Hz sensory entrainment, we further analyzed perceptual performance in those two stimulation conditions. We found that both conditions revealed no effect of phase angle at tactile stimulation onset on detection rate. While our strong effects of beta stimulation on ISPC is evidence against a lack of entrainment, we argue that the fragile effect of the ongoing oscillatory phase on tactile perception is erased by the strong effect caused by any preceding tactile stimulation. Such a masking effect occurred in our study throughout the entrainment conditions and will be discussed in the following paragraphs.

### Sensory entrainment masks subsequent tactile detection

Masking refers to the phenomenon that one sensory stimulus decreases the detectability of another when activating the sensory system simultaneously or in close temporal relationship (Verrillo, Gescheider, Calman, & Van Doren, 1983). Such an effect can also occur when the two stimuli are applied to different locations on the skin (e.g. different fingers), which has been suggested to represent inhibition effects across neurons which process projections from multiple peripheral locations (Biermann et al., 1998; Forss, Jousmäki, & Hari, 1995).

The stimulation paradigm in the present study is prone to such a masking effect, because sensory entrainment (vibrotactile stimulation) of the thumb is preceding a weak tactile stimulus to the index finger within 25 ms to 75 ms. Exploration of the estimated thresholds derived from each entrainment condition revealed clear decreases in detectability (i.e. increases in estimated thresholds) in comparison to control without preceding entrainment. This effect was stronger for high-intensity entrainment conditions compared to low, reflecting stronger masking effects. Further, this finding was independent of stimulation rhythmicity (rhythmic vs. jittered) and frequency (10 Hz vs. 20 Hz).

### Masking effect depends on strength of neural entrainment in beta

Since masking is believed to relate to the neuronal processing of sensory information, and entrainment modulates the neural activity which is thought to underlie these processes, we investigated whether a relationship between the masking effect and the gain of the sensory entrainment (i.e. the increase of ISPC from low-to high-intensity entrainment: ∆ ISPC) could be uncovered. Our analysis found a significant correlation between the entrainment gain (∆ ISPC) from low to high intensity and the behavioral masking effect (∆ TH) only for the beta band. A median-split of the data revealed that those individuals with higher neural entrainment gains had a stronger increase of detection thresholds, and therefore an increased masking effect for high-intensity 20 Hz stimulation than those with a low gain (i.e. small ∆ ISPC).

This finding demonstrates a relationship between neuronal responses to vibrotactile stimulation and perceptual processing. If entrainment effects between low- and high-intensity beta stimulation are comparable (i.e. close ISPC), masking effects on tactile detection seem to lack an increase from low-to high-intensity stimulation. The fact that we find this relationship only in beta entrainment conditions, but not in alpha or jittered, further reaffirms our assumption that the beta band is most sensitive to rhythmic tactile entrainment.

Previously, we showed that ISPC between a vibrotactile beta stimulation and the EEG response in the contralateral somatosensory area revealed characteristics of the Arnold tongue (Wälti et al., 2019). While high-intensity beta stimulation resulted in high ISPC independent from stimulation frequency, we found that ISPC derived from low-intensity beta stimulation was dependent on the distance between the stimulation frequency and endogenous beta frequency (IBF). In other words, the closer the stimulation frequency to IBF, the closer were entrainment effects between low- and high-intensity beta stimulation. Even though we did not determine IBF (because the experiment was already very long), we speculate that in the present study variations in ∆ ISPC might reflect individual differences regarding the distance from the stimulation frequency to participants’ IBF. Based on our previous results (Wälti et al., 2019), this would result in variations of entrainment effects and consequently differences in sensory processing.

### Is rhythmic sensory stimulation a feasible method to modulate sensory perception?

Whether rhythmic sensory stimulation represents a feasible method to modulate brain oscillations and human behavior, is still debated. The main advantage, compared to tACS where effects measured by electrophysiology (e.g. EEG) can be covered by strong stimulation artifacts, is the possibility to modulate human behavior and simultaneously measure underlying brain oscillations without inducing artifacts. However, as shown here, rhythmic sensory stimulation entails strong caveats when aiming to modulate sensory perception processes, which are not present in other forms of neural entrainment (e.g. tACS). We found that supra-threshold sensory stimulation entails unwanted effects (e.g. masking) on perception, which overshadow possible effects of entrained neural oscillations.

Our findings show how both of these mechanisms can affect the perception threshold of a tactile stimulus. Masking occurs regardless of stimulation frequency and rhythmicity, and appears stronger (higher thresholds) for higher stimulation intensities. Within this masking, however, neural entrainment in the beta band can have a modulatory effect on perceptual thresholds as well.

Because of large side-effects on the targeted sensory systems, the feasibility of rhythmic sensory stimulation to modulate perception remains questionable. Possible approaches to overcome this issue would be to stimulate other sensory systems (aiming for cross-modal entrainment) or presenting stimulation at a sub-threshold intensity. However, stimulation of one modality is thought to have attentional effects on other modalities (cross-modal attention; see Macaluso, Frith, & Driver, 2002). Further, entrainment of an oscillating system requires sufficient force (intensity) and would arguably not result from sub-threshold stimulation (see Thut et al., 2011). In conclusion, rhythmic sensory stimulation represents a possible but not preferable method to modulate perception in the somatosensory system when compared to other forms of neural entrainment.

## Conclusion

Our study shows that rhythmic sensory stimulation causes resonance-like effects of brain activity that are present in the EEG signal. Moreover, it can alter neuronal information processing by modulating sensory perception. Even though every supra-threshold sensory stimulation appears to affect sensory perception, the only significant brain-behavior correlation was found for entrainment in the beta band, confirming that beta activity is closely linked to somatosensory processing.

## Acknowledgments

We thank Alexandra Bürgler, Lena Salzmann, and Alexander Hess for their help with data acquisition, and Finn Rabe for methodological support. We also acknowledge the support of the Neuroscience Center Zurich (ZNZ).

## Notes

**Conflict of interest** The authors declare that the research was conducted in the absence of any commercial or financial relationships that could be construed as a potential conflict of interest.

## References

Adrian, E. D., & Yamagiwa, K. (1935). The origin of the Berger rhythm. Brain: A Journal of Neurology, 58(3), 323–351.

Ai, L., & Ro, T. (2014). The phase of prestimulus alpha oscillations affects tactile perception. J Neurophysiol, 11.(6), 1300–1307. doi: 10.1152/jn.00125.2013

Baayen, R. H., Davidson, D. J., & Bates, D. M. (2008). Mixed-effects modeling with crossed random effects for subjects and items. Journal of Memory and Language, 59(4), 390–412. doi: 10.1016/j.jml.2007.12.005

Bächinger, M., Zerbi, V., Moisa, M., Polania, R., Liu, Q., Mantini, D., … Wenderoth, N. (2017). Concurrent tACS-fMRI Reveals Causal Influence of Power Synchronized Neural Activity on Resting State fMRI Connectivity. Journal of Neuroscience, 37(18), 4766–4777. doi: 10.1523/JNEUROSCI.1756-16.2017

Bates, D., Mächler, M., Bolker, B., & Walker, S. (2015). Fitting Linear Mixed-Effects Models Usinglme4. Journal of Statistical Software, 67(1). doi: 10.18637/jss.v067.i01

Baumgarten, T. J., Schnitzler, A., & Lange, J. (2015). Beta oscillations define discrete perceptual cycles in the somatosensory domain. Proceedings of the National Academy of Sciences, 112(39), 12187–12192.

Biermann, K., Schmitz, F., Witte, O. W., Konczak, J., Freund, H. J., & Schnitzler, A. (1998). Interaction of finger representation in the human first somatosensory cortex: a neuromagnetic study. Neuroscience letters, 251(1), 13–16.

Cecere, R., Rees, G., & Romei, V. (2015). Individual differences in alpha frequency drive crossmodal illusory perception. Curr Biol, 25(2), 231–235. doi: 10.1016/j.cub.2014.11.034

Cohen, M. X. (2014). Analyzing neural time series data: theory and practice: MIT press.

Craddock, M., Poliakoff, E., El-Deredy, W., Klepousniotou, E., & Lloyd, D. M. (2017). Pre-stimulus alpha oscillations over somatosensory cortex predict tactile misperceptions. Neuropsychologia, 96, 9–18. doi: 10.1016/j.neuropsychologia.2016.12.030

Ergenoglu, T., Demiralp, T., Bayraktaroglu, Z., Ergen, M., Beydagi, H., & Uresin, Y. (2004). Alpha rhythm of the EEG modulates visual detection performance in humans. Brain Res Cogn Brain Res, 20(3), 376–383. doi: 10.1016/j.cogbrainres.2004.03.009

Forss, N., Jousmäki, V., & Hari, R. (1995). Interaction between afferent input from fingers in human somatosensory cortex. Brain Research, 685(1-2), 68–76.

Frey, J. N., Ruhnau, P., Leske, S., Siegel, M., Braun, C., & Weisz, N. (2016). The Tactile Window to Consciousness is Characterized by Frequency-Specific Integration and Segregation of the Primary Somatosensory Cortex. Sci Rep, 6, 20805. doi: 10.1038/srep20805

Gilson, R. D. (1969). Vibrotactile masking: Some spatial and temporal aspects. Perception & Psychophysics, 5(3), 176–180.

Gundlach, C., Muller, M. M., Nierhaus, T., Villringer, A., & Sehm, B. (2017). Modulation of Somatosensory Alpha Rhythm by Transcranial Alternating Current Stimulation at Mu-Frequency. Front Hum Neurosci, 11, 432. doi: 10.3389/fnhum.2017.00432

Gundlach, C., Müller, M. M., Nierhaus, T., Villringer, A., & Sehm, B. (2016). Phasic Modulation of Human Somatosensory Perception by Transcranially Applied Oscillating Currents. Brain Stimulation, 9(5), 712–719. doi: 10.1016/j.brs.2016.04.014

Haegens, S., & Zion Golumbic, E. (2018). Rhythmic facilitation of sensory processing: A critical review. Neurosci Biobehav Rev, 86, 150–165. doi: 10.1016/j.neubiorev.2017.12.002

Hanslmayr, S., Aslan, A., Staudigl, T., Klimesch, W., Herrmann, C. S., & Bauml, K. H. (2007). Prestimulus oscillations predict visual perception performance between and within subjects. Neuroimage, 37(4), 1465–1473. doi: 10.1016/j.neuroimage.2007.07.011

Hanslmayr, S., Gross, J., Klimesch, W., & Shapiro, K. L. (2011). The role of alpha oscillations in temporal attention. Brain Res Rev, 67(1-2), 331–343. doi: 10.1016/j.brainresrev.2011.04.002

Harvey, B. M., Vansteensel, M. J., Ferrier, C. H., Petridou, N., Zuiderbaan, W., Aarnoutse, E. J., … Dumoulin, S. O. (2013). Frequency specific spatial interactions in human electrocorticography: V1 alpha oscillations reflect surround suppression. Neuroimage, 65, 424–432. doi: 10.1016/j.neuroimage.2012.10.020

Helfrich, R. F., Schneider, T. R., Rach, S., Trautmann-Lengsfeld, S. A., Engel, A. K., & Herrmann, C. S. (2014). Entrainment of brain oscillations by transcranial alternating current stimulation. Curr Biol, 24(3), 333–339. doi: 10.1016/j.cub.2013.12.041

Hutcheon, B., & Yarom, Y. (2000). Resonance, oscillation and the intrinsic frequency preferences of neurons. Trends in Neuroscience, 23, 216–222.

Iemi, L., Chaumon, M., Crouzet, S. M., & Busch, N. A. (2017). Spontaneous Neural Oscillations Bias Perception by Modulating Baseline Excitability. The Journal of Neuroscience, 37(4), 807–819. doi: 10.1523/jneurosci.1432-16.2016

Jensen, O., & Mazaheri, A. (2010). Shaping functional architecture by oscillatory alpha activity: gating by inhibition. Front Hum Neurosci, 4, 186. doi: 10.3389/fnhum.2010.00186

Klimesch, W., Sauseng, P., & Hanslmayr, S. (2007). EEG alpha oscillations: the inhibition-timing hypothesis. Brain Res Rev, 53(1), 63–88. doi: 10.1016/j.brainresrev.2006.06.003

Lange, J., Oostenveld, R., & Fries, P. (2013). Reduced occipital alpha power indexes enhanced excitability rather than improved visual perception. J Neurosci, 3(7), 3212–3220. doi: 10.1523/JNEUROSCI.3755-12.2013

Limbach, K., & Corballis, P. M. (2016). Prestimulus alpha power influences response criterion in a detection task. Psychophysiology, 53(8), 1154–1164. doi: 10.1111/psyp.12666

Linkenkaer-Hansen, K., Nikulin, V. V., Palva, S., Ilmoniemi, R. J., & Palva, J. M. (2004). Prestimulus oscillations enhance psychophysical performance in humans. J Neurosci, 2(45), 10186–10190. doi: 10.1523/JNEUROSCI.2584-04.2004

Liu, Q., Ganzetti, M., Wenderoth, N., & Mantini, D. (2018). Detecting Large-Scale Brain Networks Using EEG: Impact of Electrode Density, Head Modeling and Source Localization. Front Neuroinform, 12, 4. doi: 10.3389/fninf.2018.00004

Macaluso, E., Frith, C. D., & Driver, J. (2002). Directing attention to locations and to sensory modalities: multiple levels of selective processing revealed with PET. Cerebral Cortex, 12(4), 357–368.

Mathewson, K. E., Gratton, G., Fabiani, M., Beck, D. M., & Ro, T. (2009). To see or not to see: prestimulus alpha phase predicts visual awareness. J Neurosci, 2(9), 2725–2732. doi: 10.1523/JNEUROSCI.3963-08.2009

Mathewson, K. E., Lleras, A., Beck, D. M., Fabiani, M., Ro, T., & Gratton, G. (2011). Pulsed out of awareness: EEG alpha oscillations represent a pulsed-inhibition of ongoing cortical processing. Front Psychol, 2, 99. doi: 10.3389/fpsyg.2011.00099

Mazaheri, A., & Jensen, O. (2010). Rhythmic pulsing: linking ongoing brain activity with evoked responses. Front Hum Neurosci, 4, 177. doi: 10.3389/fnhum.2010.00177

Müller, G. R., Neuper, C., & Pfurtscheller, G. (2001). “Resonance-like” frequencies of sensorimotor areas evoked by repetitive tactile stimulation. Biomedical Engineering, 46(7-8), 186–190.

Notbohm, A., Kurths, J., & Herrmann, C. S. (2016). Modification of Brain Oscillations via Rhythmic Light Stimulation Provides Evidence for Entrainment but Not for Superposition of Event-Related Responses. Front Hum Neurosci, 10, 10. doi: 10.3389/fnhum.2016.00010

Palva, S., Linkenkaer-Hansen, K., Naatanen, R., & Palva, J. M. (2005). Early neural correlates of conscious somatosensory perception. J Neurosci, 2(21), 5248–5258. doi: 10.1523/JNEUROSCI.0141-05.2005

Pfurtscheller, G., & Lopes da Silva, F. H. (1999). Event-related EEG/MEG synchronization and desynchronization: basic principles. Clinical Neurophysiology, 110, 1842–1857.

R Core Team. (2018). R: A language and environment for statistical computing. Vienna, Austria. Retrieved from https://www.r-project.org.

Rajagovindan, R., & Ding, M. (2011). From Prestimulus Alpha Oscillation to Visual-evoked Response: An Inverted-U Function and Its Attentional Modulation. Journal of Cognitive Neuroscience, 23(6), 1379–1394.

Regan, D. (1977). Steady-state evoked potentials. JOSA, 67(11), 1475–1489.

Romei, V., Brodbeck, V., Michel, C., Amedi, A., Pascual-Leone, A., & Thut, G. (2008). Spontaneous fluctuations in posterior alpha-band EEG activity reflect variability in excitability of human visual areas. Cereb Cortex, 18(9), 2010–2018. doi: 10.1093/cercor/bhm229

Romei, V., Gross, J., & Thut, G. (2010). On the role of prestimulus alpha rhythms over occipito-parietal areas in visual input regulation: correlation or causation? J Neurosci, 3(25), 8692–8697. doi: 10.1523/JNEUROSCI.0160-10.2010

Schubert, R., Haufe, S., Blankenburg, F., Villringer, A., & Curio, G. (2009). Now you’ll feel it, now you won’t: EEG rhythms predict the effectiveness of perceptual masking. Journal of Cognitive Neuroscience, 21(12), 2407–2419.

Schwab, K., Ligges, C., Jungmann, T., Hilgenfeld, B., Haueisen, J., & Witte, H. (2006). Alpha entrainment in human electroencephalogram and magnetoencephalogram recordings. Neuroreport, 17(17), 1829–1833.

Snyder, A. Z. (1992). Steady-state vibration evoked potentials: description of technique and characterization of responses. Electroencephalography and Clinical Neurophysiology/Evoked potentials Section, 84(3), 257–268.

Thut, G., Nietzel, A., Brandt, S. A., & Pascual-Leone, A. (2006). Alpha-band electroencephalographic activity over occipital cortex indexes visuospatial attention bias and predicts visual target detection. J Neurosci, 2(37), 9494–9502. doi: 10.1523/JNEUROSCI.0875-06.2006

Thut, G., Schyns, P. G., & Gross, J. (2011). Entrainment of perceptually relevant brain oscillations by non-invasive rhythmic stimulation of the human brain. Front Psychol, 2, 170. doi: 10.3389/fpsyg.2011.00170

Tobimatsu, S., Zhang, Y. M., & Kato, M. (1999). Steady-state vibration somatosensory evoked potentials: physiological characteristics and tuning function. Clinical Neurophysiology, 110(11), 1953–1958.

Tobimatsu, S., Zhang, Y. M., Suga, R., & Kato, M. (2000). Differential temporal coding of the vibratory sense in the hand and foot in man. Clinical Neurophysiology, 111(3), 398–404.

van Dijk, H., Schoffelen, J. M., Oostenveld, R., & Jensen, O. (2008). Prestimulus oscillatory activity in the alpha band predicts visual discrimination ability. J Neurosci, 2(8), 1816–1823. doi: 10.1523/JNEUROSCI.1853-07.2008

Verrillo, R. T., Gescheider, G. A., Calman, B. G., & Van Doren, C. L. (1983). Vibrotactile masking: Effects of oneand two-site stimulation. Perception & Psychophysics, 33(4), 379–387.

Vialatte, F. B., Maurice, M., Dauwels, J., & Cichocki, A. (2010). Steady-state visually evoked potentials: focus on essential paradigms and future perspectives. Prog Neurobiol, 90(4), 418–438. doi: 10.1016/j.pneurobio.2009.11.005

Wälti, M. J., Bächinger, M., Ruddy, K. L., & Wenderoth, N. (2019). Steady-state responses in the somatosensory system interact with endogenous beta activity. bioRxiv 690495. doi: 10.1101/690495

Watson, A. B., & Pelli, D. G. (1983). QUEST: A Bayesian adaptive psychometric method. Perception & Psychophysics, 33(2), 113–120.

Weisz, N., Wuhle, A., Monittola, G., Demarchi, G., Frey, J., Popov, T., & Braun, C. (2014). Prestimulus oscillatory power and connectivity patterns predispose conscious somatosensory perception. Proc Natl Acad Sci U S A, 111(4), E417–425. doi: 10.1073/pnas.1317267111

Zaehle, T., Rach, S., & Herrmann, C. S. (2010). Transcranial alternating current stimulation enhances individual alpha activity in human EEG. PLoS One, 5(11), e13766. doi: 10.1371/journal.pone.0013766

Zhang, Y., & Ding, M. (2010). Detection of a weak somatosensory stimulus: Role of the prestimulus mu rhythm and its top–down modulation. Journal of Cognitive Neuroscience, 22(2), 307–322.

Zoefel, B., Ten Oever, S., & Sack, A. T. (2018). The Involvement of Endogenous Neural Oscillations in the Processing of Rhythmic Input: More Than a Regular Repetition of Evoked Neural Responses. Front Neurosci, 12, 95. doi: 10.3389/fnins.2018.00095

